## Supplementary Material for "Neonatal brain dynamic functional connectivity in term and preterm infants and its association with early childhood neurodevelopment"

**Supplementary Table S1 | Association of global dynamic features (synchrony and metastability) with PMA and PND at scan, and effect of preterm birth using M-CRIB atlas.**

| | Term (*n* = 324) | | | | Term vs Preterm (*n* = 390) | | | |
|---|---|---|---|---|---|---|---|---|
| | PMA at scan | | PND at scan | | Term (*n* = 324) | Preterm (*n* = 66) | Cohen's *D* | *p*-value[††] |
| | t[†] | p-value[†] | t[†] | p-value[†] | [mean (S.D.)] | | | |
| Mean synchronisation | -0.173 | 0.862 | 1.370 | 0.172 | 0.53 (0.08) | 0.48 (0.08) | 0.628 | $p <$ 0.001* |
| Metastability | 0.753 | 0.448 | -2.330 | 0.019* | 0.20 (0.02) | 0.19 (0.02) | 0.480 | $p <$ 0.001* |

[†]GLM1 (including 324 term-born babies): $y \sim \beta_0 + \beta_1$PMA + $\beta_2$PND + $\beta_3$Sex + $\beta_4$Motion outliers (FD). [††]GLM2 (including 324 term-born and 66 preterm-born babies): $y \sim \beta_0 + \beta_1$Preterm-born + $\beta_2$PMA + $\beta_3$Sex + $\beta_4$Motion outliers (FD). p-values obtained with a two-sided permutation test. * Indicates results surviving FDR multiple comparison correction with $\alpha$ error at 5%). PMA: Postmenstrual age. PND: Postnatal days.

### Supplementary Figures

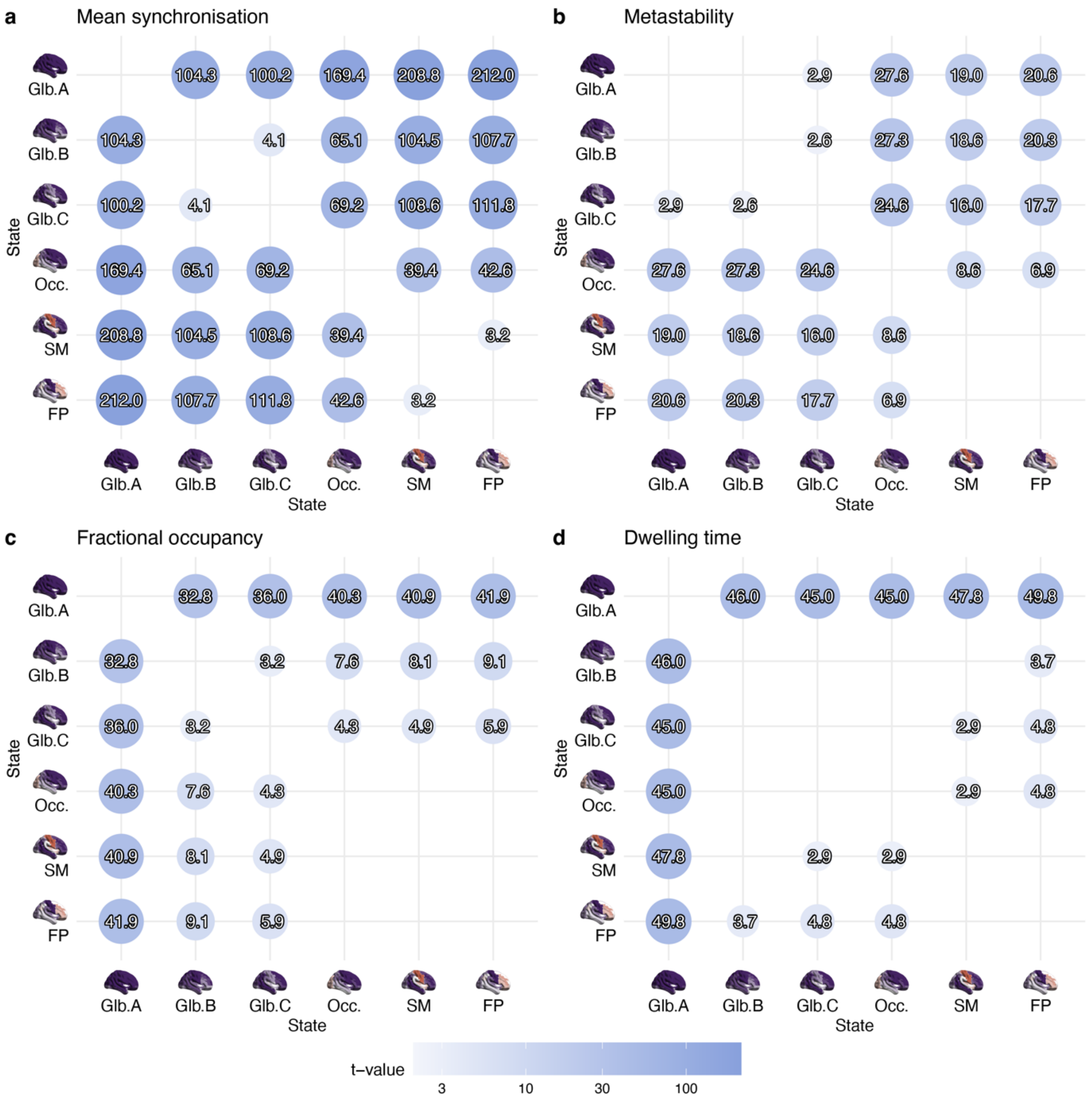


**Supplementary Figure S1 |** ANOVA two-sided post-hoc tests (*n* = 324) for a) Mean synchronisation, b) Metastability, c) fractional occupancy, and d) dwell times. Glb.: Global. Occ.: Occipital. SM: Sensorimotor. FP: Frontoparietal.

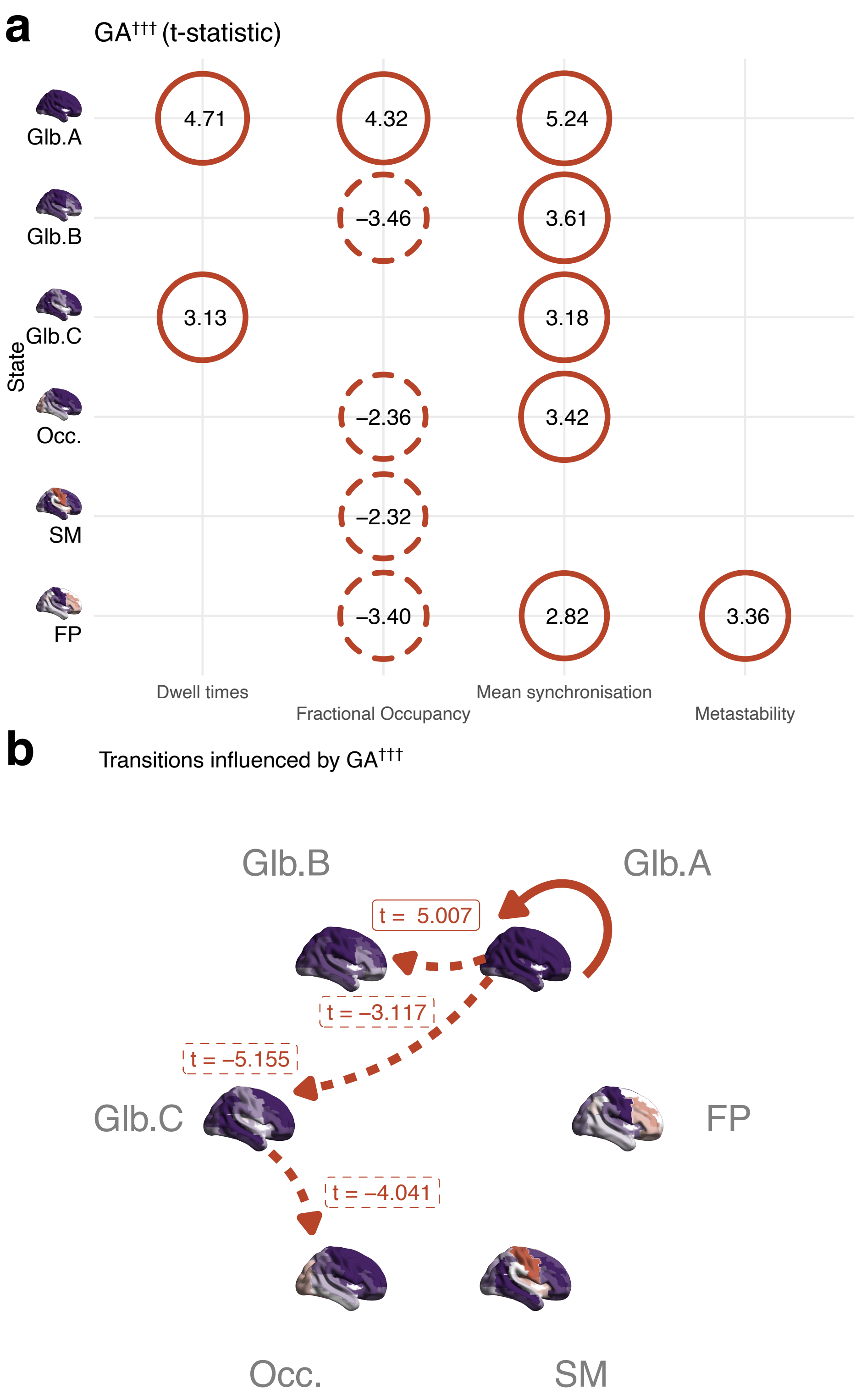


**Supplementary Figure S2 | Association of GA at birth with modular features of neonatal brain dynamics (n = 390).** (A) Summary of associations between each of the four metrics (dwell times, fractional occupancy, mean synchronisation, metastability) and GA at birth. (**B**) Association of state transitions probabilities and GA at birth. †††GLM3 (including 324 term-born and 66 preterm-born babies): $y \sim \beta_0 + \beta_1\text{GA} + \beta_2\text{PMA} + \beta_3\text{Sex} + \beta_4\text{Motion outliers (FD)}$. * p < 0.05. ** p < 0.01. *** p < 0.001. Glb.: Global. Occ.: Occipital. SM: Sensorimotor. FP: Frontoparietal. PMA: Postmenstrual age. PND: Postnatal days. GA: Gestational age.

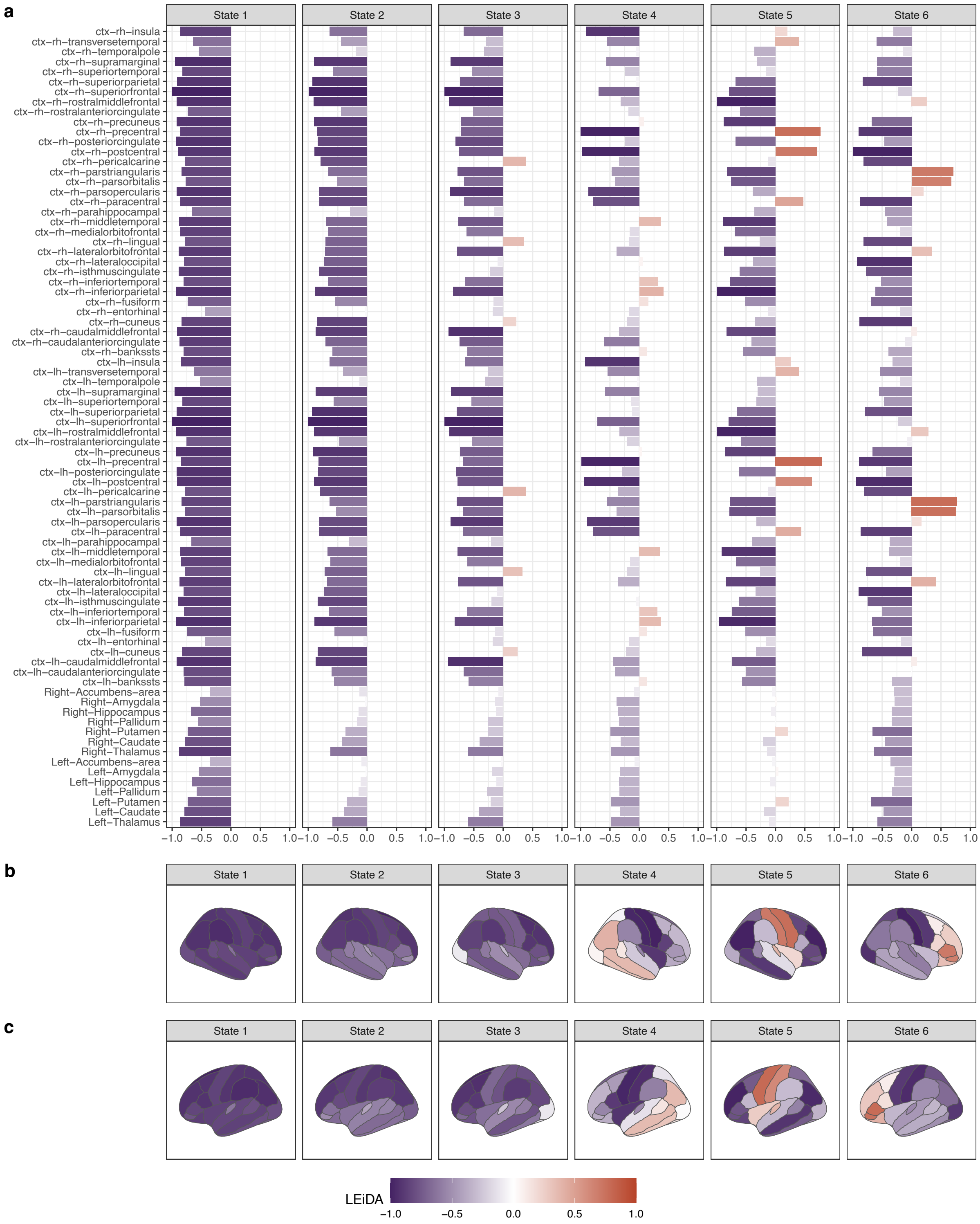


**Supplementary Figure S3 | Brain states in neonates using M-CRIB atlas.** (**a**) LEiDA vectors for each of the six brain states identified in the neonatal brain using M-CRIB parcels. (**b**) Representation of LEiDA on brain surfaces (right side view). (**c**) Representation of LEiDA on brain surfaces (left side view).

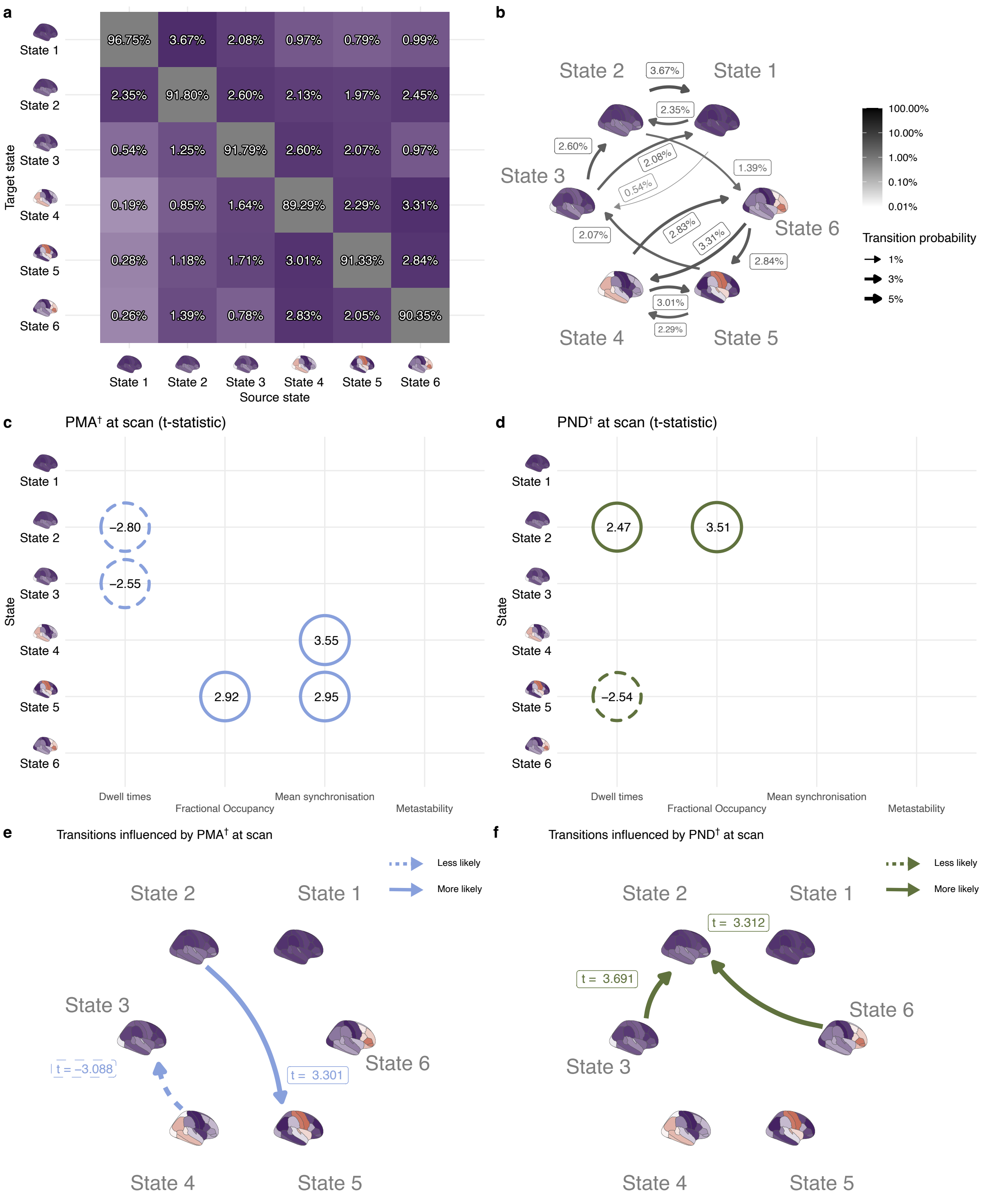


**Supplementary Figure S4 | Brain dynamics in term-born children (n = 324) (M-CRIB atlas).** (**a**) All transitions including dwelling state transitions. (**b**) Main transitions (top 12) between states excluding dwelling sequences. (**c**) Summary of brain state features significantly associated with PMA at scan. (**d**) Summary of brain state features significantly associated with PND at scan. (**e**) Summary of significant correlations between state transitions probabilities and PMA at scan. (**f**) Summary of significant correlations between state transitions probabilities and PND at scan. †GLM1 (including 324 term-born babies): $y \sim \beta_0 + \beta_1\text{PMA} + \beta_2\text{PND} + \beta_3\text{Sex} + \beta_4\text{Motion outliers (FD)}$. Values shown in **c** and **d** indicate t-statistics. All significant associations (two-sided permutation test) shown in this figure survive FDR multiple comparison correction with α error at 5%. PMA: Postmenstrual age. PND: Postnatal days.

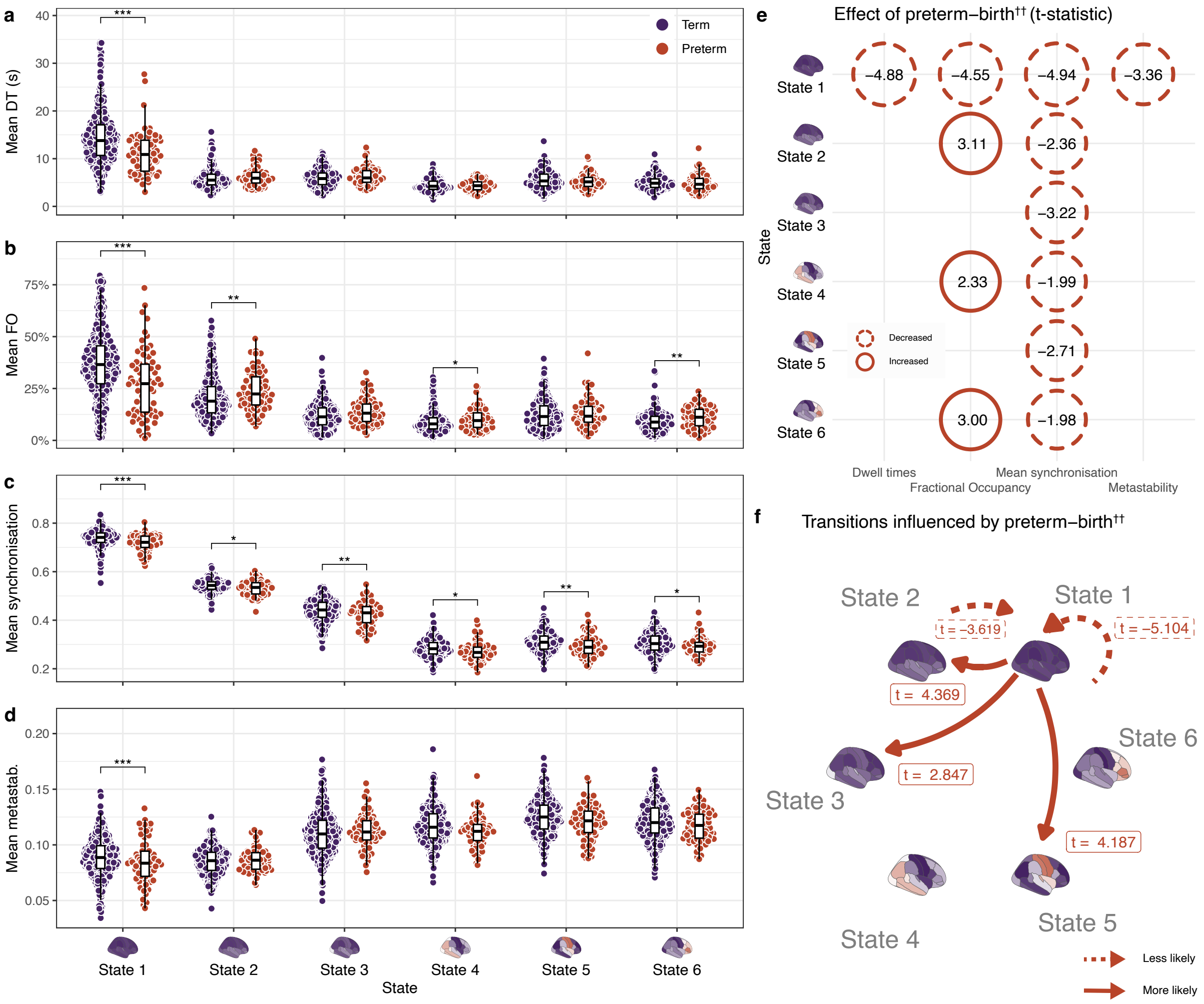


**Supplementary Figure S5 | Effect of preterm birth in brain dynamics (M-CRIB atlas).** (**a**) Mean dwell times (DT). (**b**) Mean fractional occupancy (FO). (**c**) Mean synchronisation. (**d**) Metastability. (**e**) Summary of significant associations with preterm birth (**f**) Association of state transitions probabilities and preterm birth. †GLM1 (324 term-born babies): $y \sim \beta_0 + \beta_1 PMA + \beta_2 PND + \beta_3 Sex + \beta_4 Motion\ outliers\ (FD)$. ††GLM2 (324 term-born and 66 preterm-born babies): $y \sim \beta_0 + \beta_1 Preterm\text{-}born + \beta_2 PMA + \beta_3 Sex + \beta_4 Motion\ outliers\ (FD)$. * $p < 0.05$. ** $p < 0.01$. *** $p < 0.001$ obtained with a two-sided permutation test. Values shown in **e** indicate t-statistics. Boxplots showing 0th, 25th, 50th, 75th and 100th centiles. Outliers defined when value larger than 1.5*IQR + 75th centile. All significant associations highlighted survive FDR multiple comparison correction with α error at 5%. PMA: Postmenstrual age. PND: Postnatal days. DT: Dwell time. FO: Fractional occupancy. Metastab.: Metastability.

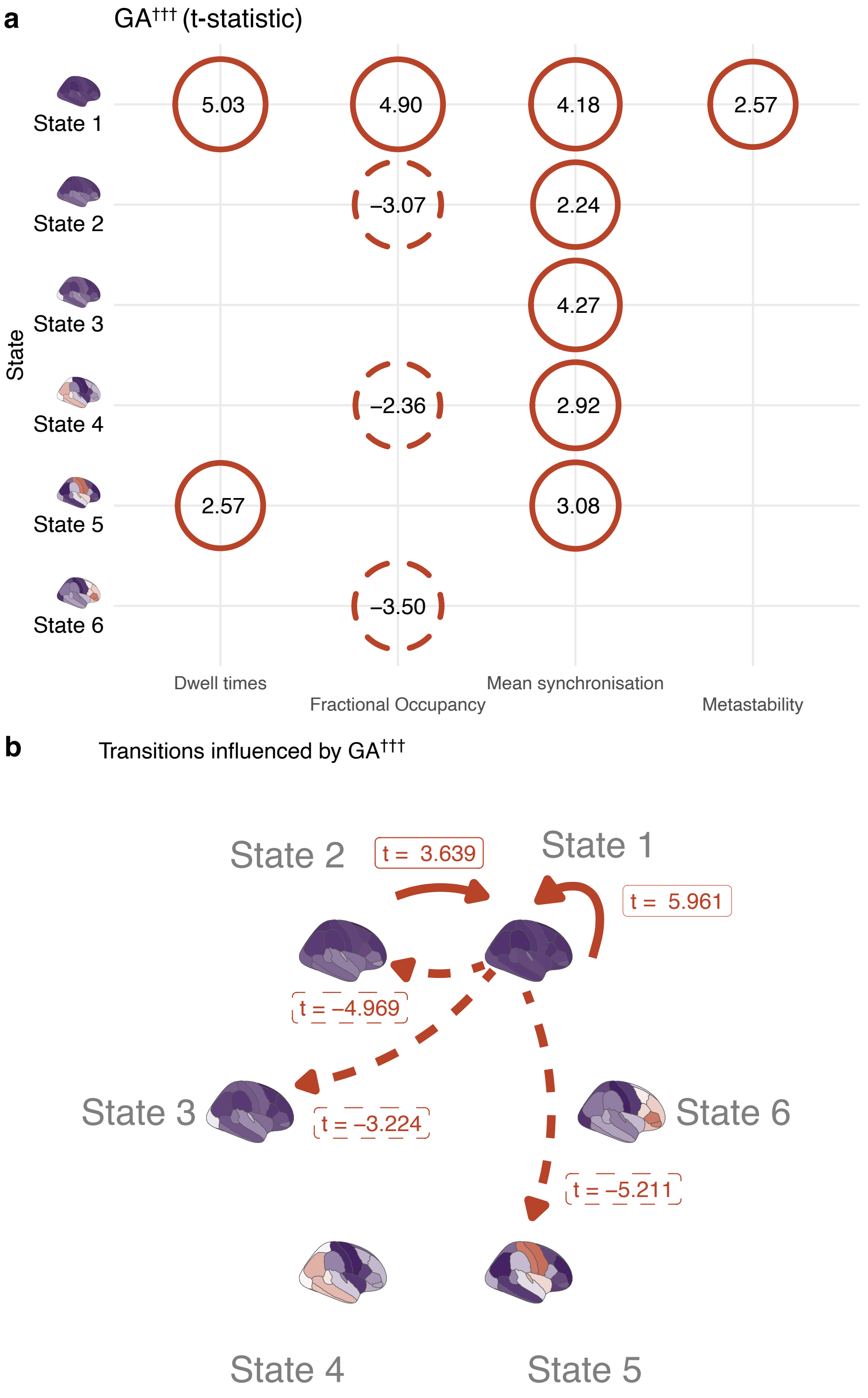


**Supplementary Figure S6 | Association of GA at birth with modular features of neonatal brain dynamics (M-CRIB atlas).** (**A**) Summary of associations between each of the four metrics (dwell times, fractional occupancy, mean synchronisation, metastability) and GA at birth. (**B**) Association of state transitions probabilities and GA at birth. †††GLM3 (including 324 term-born and 66 preterm-born babies): $y \sim \beta_0 + \beta_1\text{GA} + \beta_2\text{PMA} + \beta_3\text{Sex} + \beta_4\text{Motion outliers (FD)}$. * $p < 0.05$. ** $p < 0.01$. *** $p < 0.001$. Occ.: Occipital. SM: Sensorimotor. FP: Frontoparietal. PMA: Postmenstrual age. GA: Gestational age.

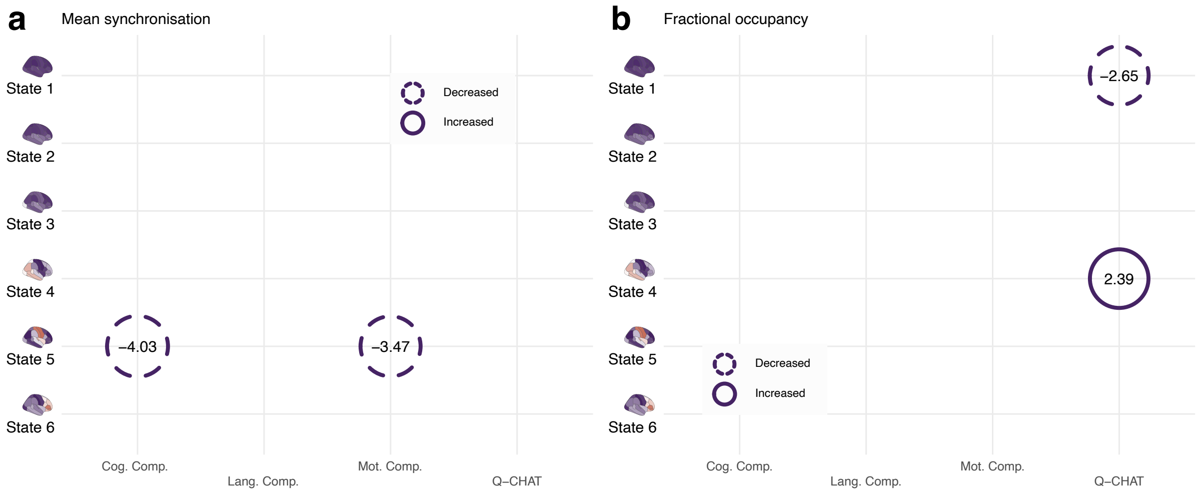


**Supplementary Figure S7 | Summary of associations of brain state features with neurodevelopmental outcomes at 18 months corrected age (M-CRIB atlas).** Association of average (**a**) mean synchronisation and (**b**) fractional occupancy in each of the six defined brain states during perinatal period with cognitive (Cog. Comp), language (Lang. Comp.) and motor (Mot. Comp) Bayley-III composite scores and Q-CHAT scores. GLM3 (257 term-born and 48 preterm-born babies for Bayley-III; and 254 term-born and 46 preterm-born babies for Q-CHAT): $y \sim \beta_0 + \beta_1$GA + $\beta_2$PMA + $\beta_3$Sex + $\beta_4$Motion outliers (FD) + $\beta_5$[Corrected age at assessment] + $\beta_6$[Assessed component] + $\beta_7$[Index of multiple deprivation]. Values shown in in both panels indicate t-statistics. All significant associations (two-sided permutation test) highlighted survive FDR multiple comparison correction with $\alpha$ error at 5%. PMA: Postmenstrual age. GA: Gestational age. Cog.: Cognitive. Lang.: Language. Mot.: Motor. Comp.: Component. Q-CHAT: Quantitative Checklist for Autism in Toddlers.

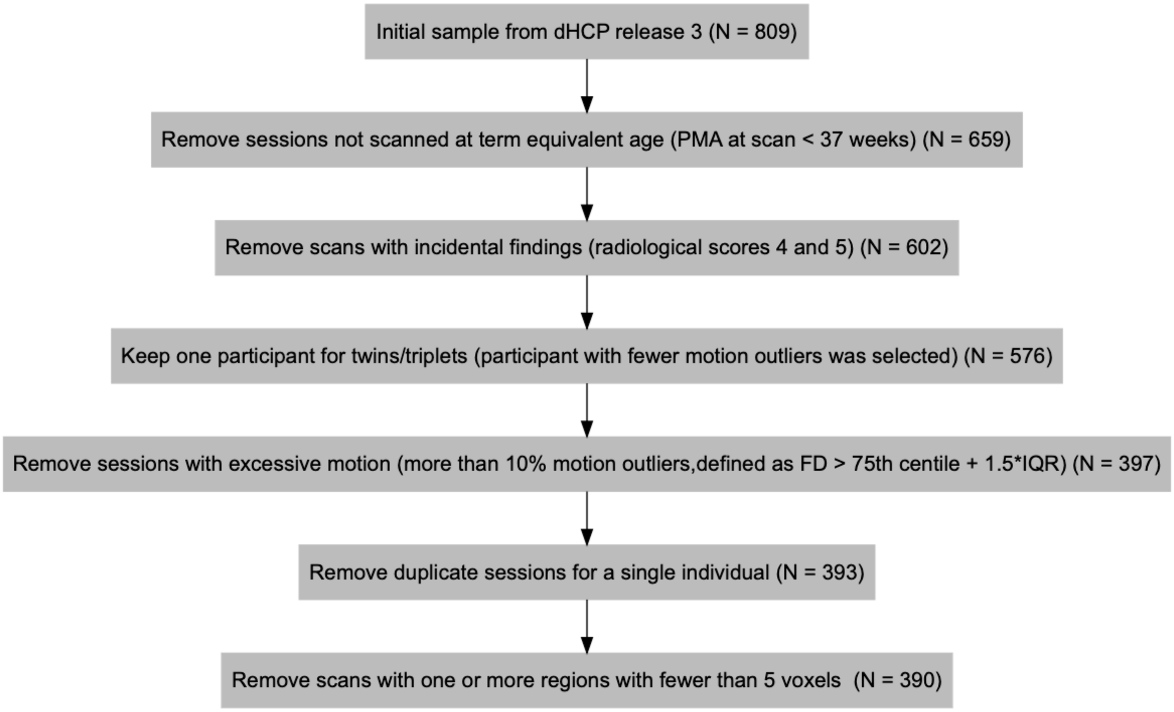


**Supplementary Figure S8.** Exclusion criteria flowchart.

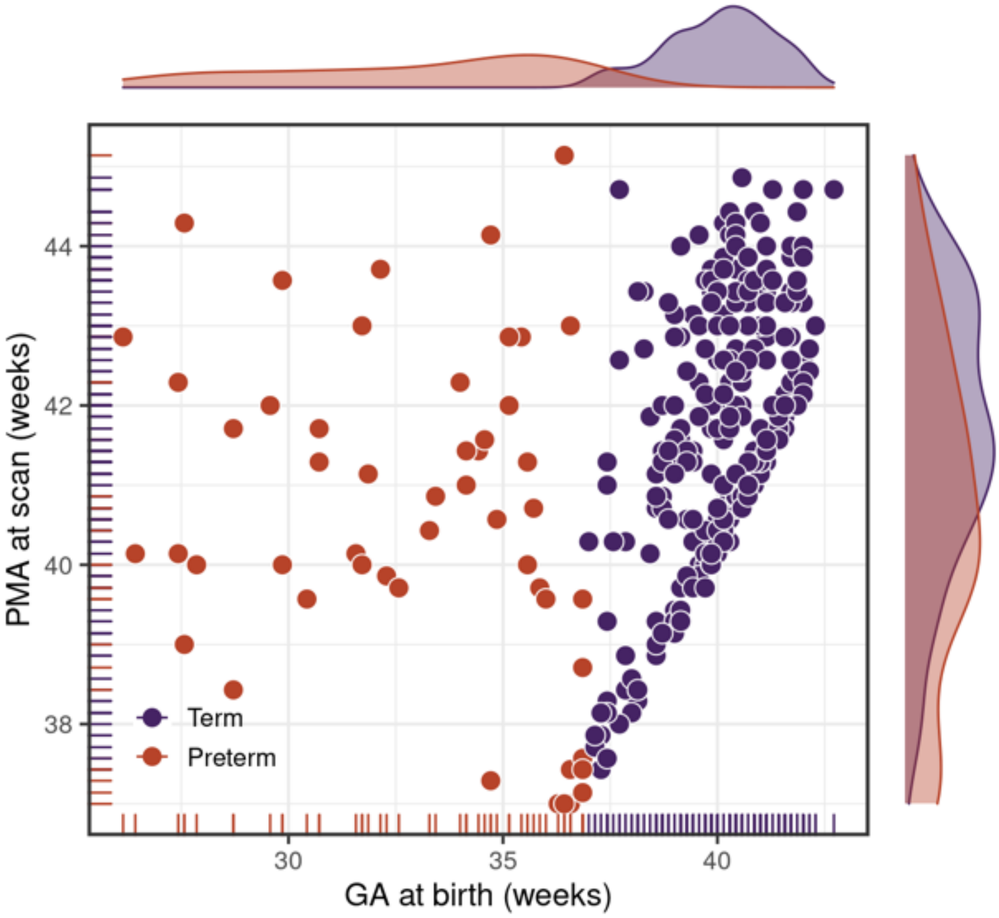


**Supplementary Figure S9.** Distribution of gestational age (GA) at birth and postmenstrual age (PMA) at scan for the participants included in this study.

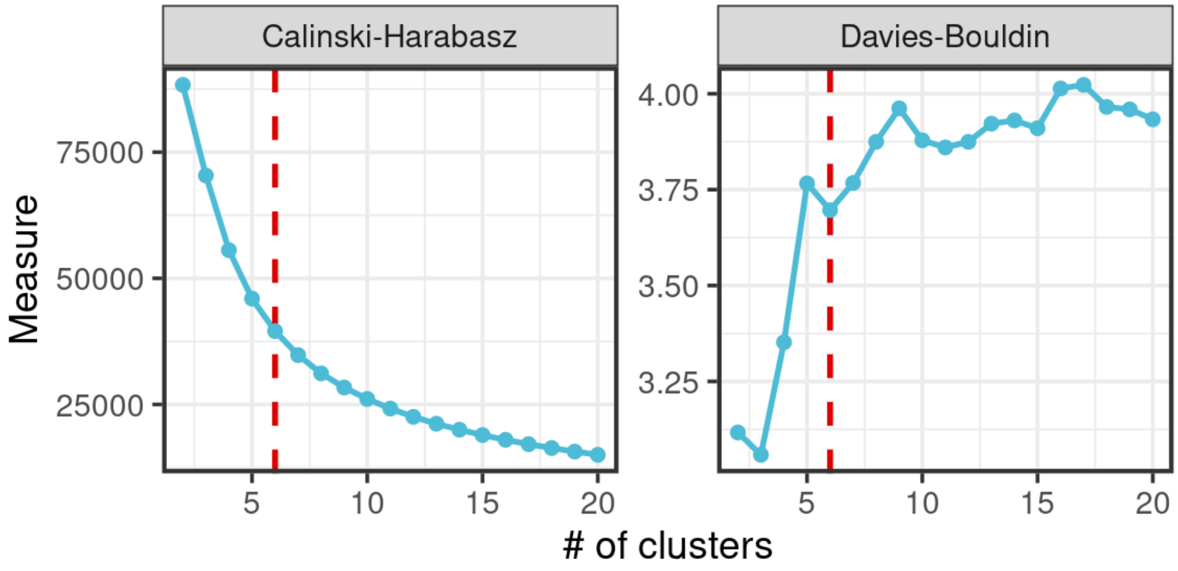


**Supplementary Figure S10.** Ideal number of clusters according to Calinski-Harabasz and Davies-Bouldin methods.

### Supplementary Results: Association of neonatal brain regional brain dynamics with PMA at scan and GA at birth using an alternative brain parcellation (M-CRIB atlas)

The six brain states obtained with the alternative M-CRIB neonatal atlas were compatible with the ones obtained for the AAL atlas (Supplementary Figure S3), with three global synchronisation states and three more regionally constrained. States 1, 2 and 3 obtained with the M-CRIB atlas were consistent with global synchronisation states obtained with the AAL atlas. The M-CRIB's State 4 featured some of the structures present in Occipital state, State 5 featured similar structures to the sensorimotor state, and State 6 was concordant with FP state obtained from AAL.

In term-born babies, PMA at scan is associated with increased mean synchronisation in State 4 ($t$ = 3.6; $p$ < 0.001) and State 5 ($t$ = 2.9; $p$ = 0.003); and fractional occupancy in State 5 ($t$ = 2.9; $p$ = 0.002). PMA is also associated with reduced dwell times for States 2 ($t$ = -2.8; $p$ = 0.004) and 3 ($t$ = -2.6; $p$ = 0.012). Transitions from State 2 to State 5 increased ($t$ = 3.3; $p$ = 0.001) and transitions from State 4 to State 3 decreased ($t$ = -3.1; $p$ = 0.003) with PMA. Also, in term-born babies, PND is associated with increases in dwell times ($t$ = 2.5; $p$ = 0.012) and fractional occupancy ($t$ = 3.5; $p$ < 0.001; and reduced dwell times for State 5 ($t$ = -2.5; $p$ = 0.012). Transitions from States 3 and 6 to State 2 also increased with PND ($t$ = 3.7; $p$ < 0.001 and $t$ = 3.3; $p$ = 0.002, respectively). SM and State 5 exhibit compatible associations with PMA and PND (Supplementary Figure S4).

Preterm-birth is associated with reduced dwell times for State 1 ($t$ = -4.9; $p$ < 0.001); reduced fractional occupancy for State 1 ($t$ = -4.5; $p$ < 0.001); reduced mean synchronisation for the States 1 ($t$ = -4.9; $p$ < 0.001), 2 ($t$ = -2.4; $p$ = 0.018), 3 ($t$ = -3.2; $p$ = 0.002), 4 ($t$ = -2.0; $p$ = 0.047), 5 ($t$ = -2.7; $p$ = 0.007), and 6 ($t$ = -2.0; $p$ < 0.05); and reduced metastability for Stage 1 ($t$ = -3.0; $p$ = 0.003). We have also found an association with increased fractional occupancy for States 2 ($t$ = 3.1; $p$ = 0.002), 4 ($t$ = 2.3; $p$ = 0.019), and 6 ($t$ = 3.0; $p$ = 0.003). Transitions from State 2 to State 1 ($t$ = -3.6; $p$ < 0.001) and dwelling sequences of State 1 ($t$ = -5.1; $p$ < 0.001) are reduced in preterm-birth. Transitions from State 1 to State 2 ($t$ = 4.4; $p$ < 0.001); State 1 to State 3 ($t$ = 2.8; $p$ = 0.005); and State 1 to State 5 ($t$ = 4.2; $p$ < 0.001) increased with preterm-birth (Supplementary Figure S5 and Supplementary Figure S6).

Developmental outcomes in cognitive ($t$ = -4.0; $p$ < 0.001) and motor ($t$ = -3.5; $p$ < 0.001) components of Bayley-III show negative associations with mean synchronisation for State 5. Moreover, higher Q-CHAT scores were associated with reduced fractional occupancy for State 1 ($t$ = -2.7; $p$ = 0.008) and increased fractional occupancy ($t$ = 2.4; $p$ = 0.017), see Supplementary Figure S7 for a summary. The results for Mean synchronisation for State 5 are compatible with those shown in AAL atlas (SM state), as well as fractional occupancy and Q-CHAT for State 1.
